# Chikungunya virus glycoproteins transform macrophages into productive viral dissemination vessels

**DOI:** 10.1101/2023.05.29.542714

**Authors:** Zhenlan Yao, Sangeetha Ramachandran, Serina Huang, Yasaman Jami-Alahmadi, James A. Wohlschlegel, Melody M.H. Li

## Abstract

Despite their role as innate sentinels, macrophages are cellular reservoirs for chikungunya virus (CHIKV), a highly pathogenic arthropod-borne alphavirus that has caused unprecedented epidemics worldwide. Here, we took interdisciplinary approaches to elucidate the CHIKV determinants that subvert macrophages into virion dissemination vessels. Through comparative infection using chimeric alphaviruses and evolutionary selection analyses, we discovered for the first time that CHIKV glycoproteins E2 and E1 coordinate efficient virion production in macrophages with the domains involved under positive selection. We performed proteomics on CHIKV-infected macrophages to identify cellular proteins interacting with the precursor and/or mature forms of viral glycoproteins. We uncovered two E1-binding proteins, signal peptidase complex subunit 3 (SPCS3) and eukaryotic translation initiation factor 3 (eIF3k), with novel inhibitory activities against CHIKV production. These results highlight how CHIKV E2 and E1 have been evolutionarily selected for viral dissemination likely through counteracting host restriction factors, making them attractive targets for therapeutic intervention.

## Main

Macrophages are phagocytic innate immune cells with critical functions in first-line defense against virus infection, inflammation, and priming of the adaptive immune system^1^. The sensing of viral infection by pattern recognition receptors in macrophages rapidly establishes an antiviral state through activation of the interferon (IFN) response^2^. However, some viruses such as highly pathogenic avian influenza H5N1 viruses can breach this antiviral immunity^3–5^, highlighting productive macrophage infection as an important determinant for viral virulence. Moreover, in individuals infected with human immunodeficiency virus (HIV)^6, 7^, macrophages are potential reservoirs for rebound viremia upon cessation of antiretroviral therapy^6, 8^. Therefore, targeting viral infection of macrophages is an attractive therapeutic strategy for virus eradication.

Chikungunya virus (CHIKV) is a highly pathogenic arthropod-borne alphavirus that has expanded worldwide with emerging lineages in recent decades^9, 10^. The unprecedented outbreaks from the Indian Ocean islands to Southeast Asia were caused by the novel CHIKV Indian Ocean lineage (IOL), characterized primarily by the E1-A226V mutation that adapted the virus from its principal vector *Aedes aegypti* to *Aedes albopictus*^11–13^. Although CHIKV infection is typically cleared in a few days, a significant percentage of individuals develops incapacitating arthralgia for up to 20 months^14–16^. Interestingly, CHIKV RNA and proteins persist in monocyte derived macrophages (MDMs) in spleen or synovial tissue for months in macaques and humans suffering from chronic arthralgia^17–19^. These studies propose a role for macrophages as a cellular reservoir for CHIKV persistence and a niche for inflammation that is recurrently activated by viral components^17, 20^. However, it is not clear what mechanism drives CHIKV persistence and whether this pathogenic role of macrophages is found in all arthritogenic alphavirus infections.

In contrast, o’nyong’nyong virus (ONNV), an arthritogenic alphavirus that shares the most genetic identity with CHIKV, is confined to periodic outbreaks in Africa^9, 21^. ONNV causes similar symptoms in humans, but is less virulent in mouse models, requiring a higher dose than CHIKV to reach the same level of mortality^22^. The evolutionary similarities yet epidemiological differences make CHIKV-ONNV chimeras excellent molecular tools for probing viral determinants for host adaptation. However, these studies so far have mostly focused on their differential uses of mosquito vectors, such as transmission of ONNV by *Anopheles gambiae*^23, 24^, while little is known about the molecular mechanisms underlying infection of human cells relevant for viral dissemination, such as macrophages.

Viral infection is mostly abortive in macrophages as host restriction factors either basally expressed or amplified by the IFN response suppress specific viral life cycle stages^25^. Even though CHIKV replication is active in human MDMs, it is more restricted in MDMs than in epithelial cells and fibroblasts^26^, suggesting viral suppression by macrophage restriction factors. Host antiviral immunity can impose evolutionary selective pressures on viral proteins, propelling viruses to evade or antagonize these blockades, such as the arms race between myeloid-cell-specific SAMHD1 and HIV-2 Vpx^27–29^. This prompted us to question how and to what extent the evolutionary pressure brought on by virus-host arms race has selected for increased CHIKV survival in human macrophages.

Here, we found that human primary monocyte and THP-1 derived macrophage infection with CHIKV (vaccine strain 181/clone 25) is much more efficient than that of ONNV at a step following genome replication. By utilizing a repertoire of CHIKV-ONNV chimeras, we mapped the viral determinant for efficient virion production in macrophages to the CHIKV E2 and E1 glycoproteins. Interestingly, evolutionary analysis of 397 CHIKV structural polyprotein sequences isolated from infected individuals uncovered signatures of positive selection mostly in E2 and E1 proteins. Mutating two of the positively selected residues in CHIKV to the homologous ones in ONNV (E2-V460L, E1-V1029I) attenuates virion production in 293T and BHK-J cells while the E1-V1029I mutation completely abolishes virion production in macrophages. We further performed affinity purification-mass spectrometry (AP-MS) on macrophage interactors of CHIKV glycoproteins. We discovered that E1 consistently interacts with signal peptidase complex subunit 3 (SPCS3) and eukaryotic translation initiation factor 3 (eIF3k), which block CHIKV production in macrophages. Taken together, we found that, in addition to their role in viral entry, CHIKV glycoproteins may interfere with cellular restrictions to facilitate virion dissemination in macrophages.

## Results

### CHIKV infects human macrophages more efficiently than other arthritogenic alphaviruses

To evaluate the susceptibility of macrophages to different arthritogenic alphaviruses, we infected human primary monocyte derived macrophages with EGFP-expressing Sindbis virus (SINV), Ross River virus (RRV), ONNV, and CHIKV, and quantified infection levels at 24 hours post-infection (h.p.i.) by flow cytometry (Figure.1a). Despite generally low infection rates with these alphaviruses (<1%), we observed a small percentage (0.76%) of macrophages highly infected with CHIKV, according to intracellular EGFP expression that spans 3 logs. We then compared growth kinetics of CHIKV and its closest relative, ONNV, in infected human monocytic cell line THP-1 derived macrophages by quantifying virion production in the supernatant (Figure.1b). We found that CHIKV produces 2-3 logs higher titers than ONNV throughout the infection time-course (up to 1.05×10^7^ pfu/ml for CHIKV compared to 3.75×10^4^ pfu/ml for ONNV), with the titers of both viruses peaking at 24 h.p.i. These results suggest that the small number of CHIKV-infected macrophages is extremely efficient at producing viral progeny.

**Figure 1.**
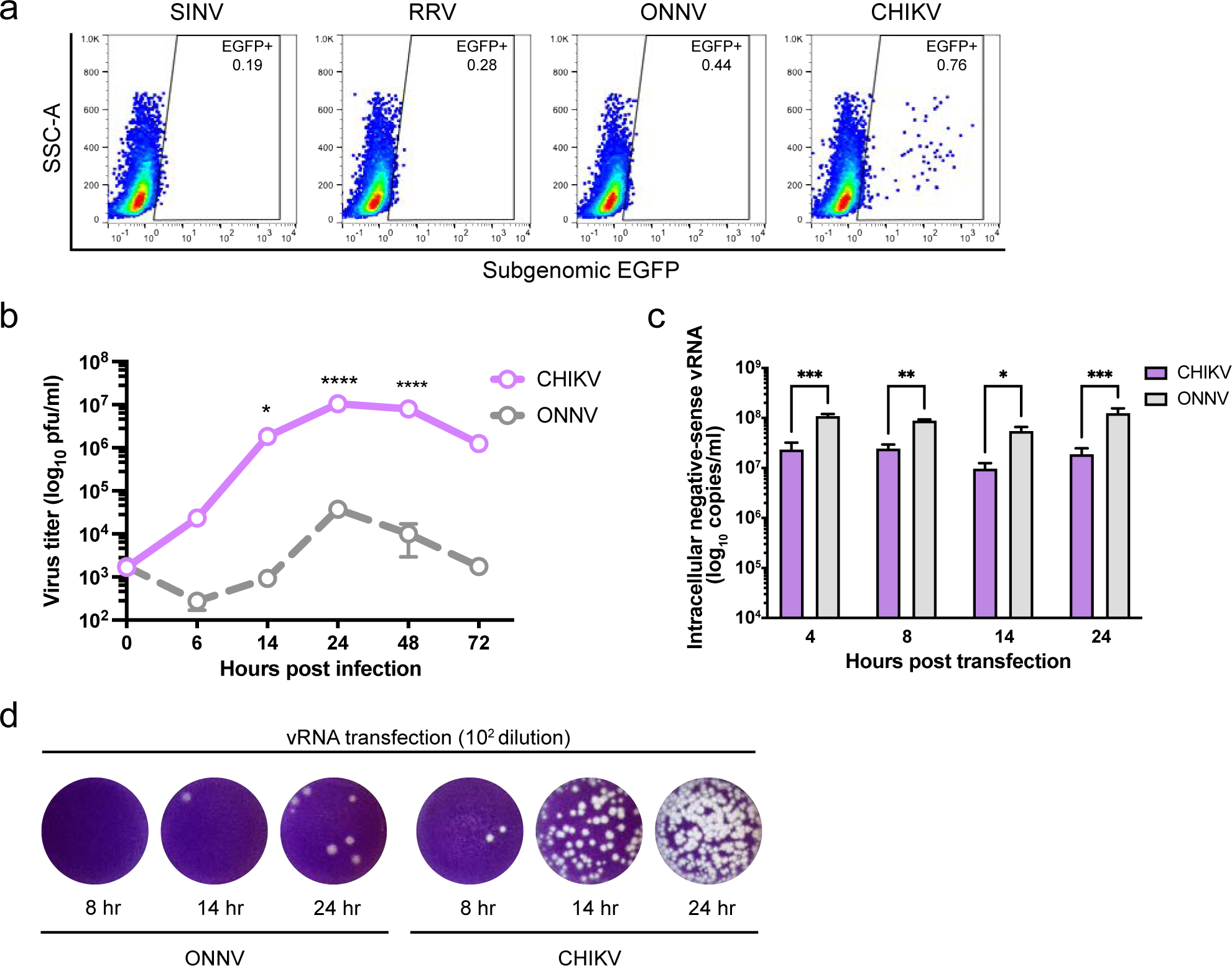
Efficient CHIKV infection in human macrophages depends on high level of virion production. **a,** Human peripheral monocyte derived macrophages were infected with EGFP-labeled alphaviruses (SINV TE/5’2J/GFP, RRV strain T48, ONNV strain SG650, and CHIKV vaccine strain 181/clone 25) at MOI of 5 for 24 h. Levels of infection with different alphaviruses were determined by percent EGFP-positive cells evaluated by flow cytometry. Data are representative of 2 independent experiments performed in biological duplicates. **b,** THP-1 derived macrophages were infected with CHIKV 181/clone 25 or ONNV SG650 at MOI 5. Titration of supernatant virus samples was performed at 0, 6, 14, 24, 48, and 72 h.p.i by plaque assay on BHK-21 cells. Data are representative of 2 independent experiments. Mean values of biological duplicates were plotted with SD. Asterisks indicate statistically significant differences as compared to ONNV (Two-way ANOVA and Šidák’s multiple comparisons test: *, p≤0.05; ****, p≤0.0001). **c,** Levels of intracellular (-) vRNAs, the viral replication intermediate, at 4, 8, 14, and 24 h post transfection of THP-1 derived macrophages with CHIKV 181/clone 25 or ONNV SG650 were quantified through RT-qPCR with specific TaqMan probes. Data are representative of 2 independent experiments. Mean values of biological duplicates measured in technical duplicates were plotted with SD (Two-way ANOVA and Šidák’s multiple comparisons test: *, p≤0.05; **, p≤0.01; ***, p≤0.001). **d,** CHIKV and ONNV titers of supernatant samples collected from transfected THP-1 derived macrophages in Figure.1c were determined by plaque assay. Representative plaques of CHIKV and ONNV from 2 independent experiments (1:100 dilution) are shown.

We asked whether the high level of CHIKV production is achieved by enhanced viral replication in macrophages. To bypass viral entry, we directly transfected in vitro transcribed genomic viral RNAs (vRNAs) of CHIKV and ONNV into THP-1-derived macrophages (Figure.1c). We measured intracellular negative-sense viral RNA ((-) vRNA), the replication intermediate, by TaqMan RT-qPCR assays. To our surprise, the (-) vRNA levels of ONNV are significantly higher than those of CHIKV following vRNA transfection, suggesting CHIKV infection is enhanced at a step after genome replication in macrophages (Figure.1c). Nevertheless, virion production of CHIKV is dramatically more robust than that of ONNV and could be detected as early as 8 hours post-transfection (h.p.t.) (Figure.1d). Taken together, human macrophage infection with CHIKV drives more superior virion production than that with ONNV.

### CHIKV E2 and E1 synergize to mediate efficient virion production in THP-1 derived human macrophages

To identify the viral determinants for CHIKV infection of human macrophages, we constructed several CHIKV-ONNV chimeras (Figure.2a) and assessed their infection levels in THP-1 derived macrophages, compared to parental CHIKV and ONNV. Alphaviruses express 4 non-structural proteins (nsP1-4) for viral replication, and 5 structural proteins from a subgenomic mRNA (capsid, E3, E2, 6K/TF, E1) for viral particle assembly and host cell entry^30^. These proteins are proteolytically processed from the non-structural and structural polyproteins. Given the genome organization, we generated Chimera I that contains ONNV nsP1 to capsid in a CHIKV backbone, and Chimera III that contains CHIKV nsP1 to capsid in an ONNV backbone. To account for potential discrepancies associated with mismatched subgenomic promoters located at the 3’ end of nsP4 and structural proteins, we also generated Chimeras II and IV, where the swapping of viral genes starts with the subgenomic promoters in CHIKV and ONNV nsP4. We found comparable levels of virion production of Chimeras I and II as CHIKV in the supernatant of infected macrophages, while Chimeras III and IV recapitulate ONNV production (Figure.2b). These data demonstrate that the viral determinants for effective macrophage infection lie in the CHIKV E3-E2-6K-E1 structural polyprotein region.

**Figure 2.**
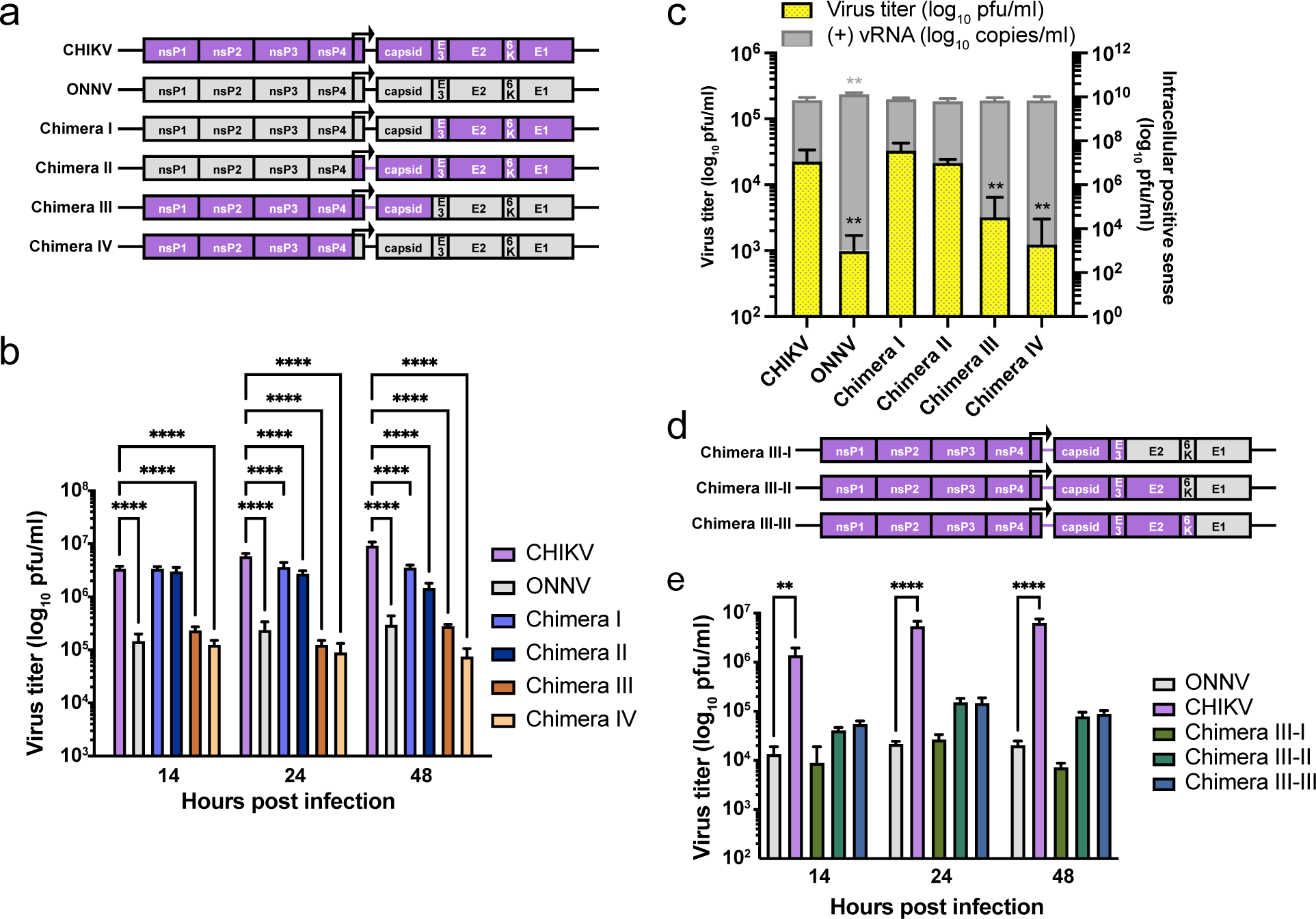
Viral glycoproteins are determinants for macrophage tropism of CHIKV. **a,** Schematic representation of CHIKV, ONNV, Chimera I, II, III, and IV. These chimeras consist of genomes from CHIKV vaccine strain 181/clone 25 and ONNV SG650 in different ratios: Chimera I contains ONNV genome from nsP1 to capsid and CHIKV genome from E3 to E1. Chimera II contains ONNV genome from nsP1 to the region prior to the subgenomic promoter in nsP4 and CHIKV genome from the subgenomic promoter to E1. Chimera III contains CHIKV genome from nsP1 to capsid and ONNV genome from E3 to E1. Chimera IV contains CHIKV genome from nsP1 to the region prior to the subgenomic promoter in nsP4 and ONNV genome from the subgenomic promoter to E1. **b,** Titration of supernatant samples from THP-1 derived macrophages infected with CHIKV 181/clone 25, ONNV SG650, and 4 chimeras (I, II, III, IV). The macrophages were inoculated with the virus at MOI 5, and the supernatant samples were collected at 14, 24, and 48 h.p.i for plaque assay analysis. Data are representative of 3 independent experiments. Mean values of biological duplicates measured in technical duplicates were plotted with SD. Asterisks indicate statistically significant differences as compared to CHIKV (Two-way ANOVA and Dunnett’s multiple comparisons test: ****, p≤0.0001). **c,** THP-1 derived macrophages were transfected with 0.5 μg RNA of CHIKV 181/clone 25, ONNV SG650, or chimeras (I, II, III, IV). Virion production was determined by intracellular (+) vRNA transcript levels and secreted infectious particle titers through RT-qPCR and plaque assay, respectively. Data are representative of 2 independent experiments. Mean values of biological duplicates measured in technical duplicates were plotted with SD. Asterisks indicate statistically significant differences as compared to CHIKV (Two-way ANOVA and Dunnett’s multiple comparisons test: **, p≤0.01). **d,** Schematic representation of Chimera III-I, III-II, and III-III. Chimera III-I contains CHIKV genome from nsP1 to E3 and ONNV genome from E2 to E1. Chimera III-II contains CHIKV genome from nsP1 to E2 and ONNV genome from 6K to E1. Chimera III-III contains CHIKV genome from nsP1 to 6K and ONNV E1. **e,** Titration of supernatant samples from THP-1 derived macrophages infected with CHIKV 181/clone 25, ONNV SG650, or chimeras (III-I, III-II, III-III) for 14, 24, and 48 h. The infection conditions and virus titer assessments were performed as previously described in Figure. 2b. Data are representative of 2 independent experiments. Mean values of biological duplicates measured in technical duplicates were plotted with SD. Asterisks indicate statistically significant differences as compared to ONNV (Two-way ANOVA and Dunnett’s multiple comparisons test: **, p≤0.01; ****, p≤0.0001).

To investigate the role of CHIKV structural proteins in virion production, we transfected vRNAs of CHIKV, ONNV, and Chimeras I-IV into THP-1 derived macrophages to bypass viral entry. We compared viral replication and production among the transfected cells at 24 h.p.t. based on intracellular positive-sense viral RNA ((+) vRNA) levels and supernatant titers (Figure.2c). Consistent with Figure.2b, transfection of viral genomes without CHIKV E3-E2-6K-E1 (ONNV, Chimera III, and Chimera IV) led to significantly lower levels of virion production.

To further narrow down the viral determinants for CHIKV infection in macrophages, we constructed three additional chimeras in the context of Chimera III to include CHIKV E3 (Chimera III-I), E3-E2 (Chimera III-II), or E3-E2-6K (Chimera III-III) (Figure.2d). Upon macrophage infection with CHIKV, ONNV, and the chimeras, we found that only Chimera III-II and Chimera III-III, both possessing CHIKV E2, partially enhance virion production at 24 and 48 h.p.i. although not significantly (Figure.2e), suggesting that E2 alone is not sufficient. Chimera III-III with all the CHIKV structural proteins except E1 fails to fully rescue virion production in macrophages. Taken together, this supports the involvement of both CHIKV E2 and E1 in virion production.

To pinpoint the impact of CHIKV E2 and E1 on virion production, we generated three chimeras in the ONNV backbone with CHIKV replacement of E2 (ONNV/CHIKV E2), E1 (ONNV/CHIKV E1), or E2 and E1 (ONNV/CHIKV E2+E1) (Figure.3a). Neither single replacement of CHIKV E2 nor E1 rescues ONNV infection of macrophages to comparable levels as CHIKV (Figure.3b. Surprisingly, macrophage infection with ONNV/CHIKV E1 is more attenuated than that with ONNV, potentially due to incompatible heterodimer formation between ONNV E2 and CHIKV E1. In contrast, simultaneous replacement of E2 and E1 with CHIKV homologs (ONNV/CHIKV E2+E1) increased the supernatant titers to levels even higher than those of CHIKV. We then transfected vRNAs into macrophages to evaluate viral replication and production (Figure.3c). All of the transfected vRNAs launched productive viral replication in macrophages; however, only the transfection of ONNV/CHIKV E2+E1 RNA led to significantly enhanced virion production, albeit at levels lower than those for transfection of CHIKV RNA (Figure.3c). These results highlight the requirement of both CHIKV E2 and E1 for efficient virion production in macrophages.

**Figure 3.**
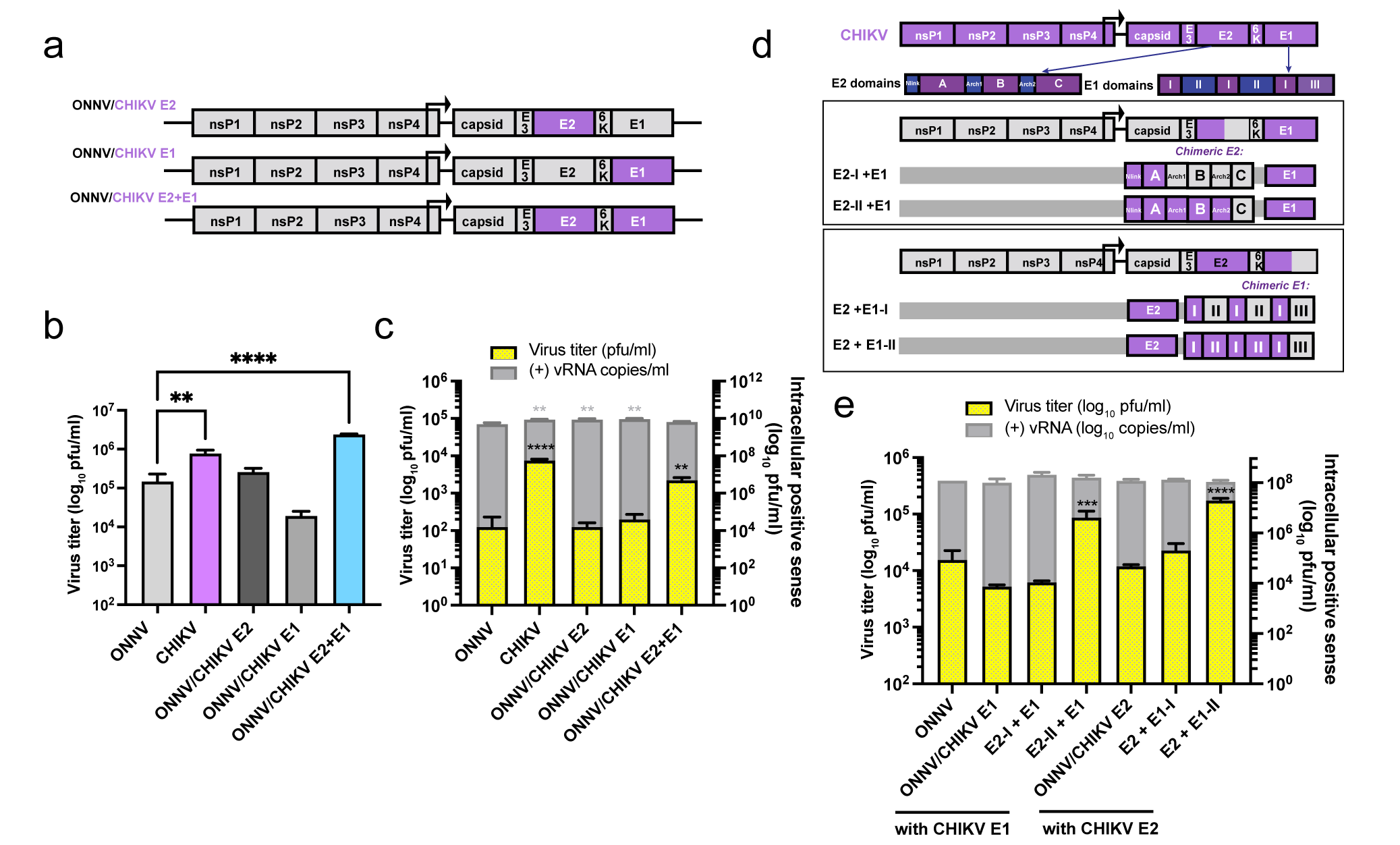
CHIKV E2 and E1 contribute to efficient virion production in human macrophages. **a,** Schematic representation of chimera ONNV/CHIKV E2, ONNV/CHIKV E1 and ONNV/CHIKV E2+E1. These three chimeric viruses were built on ONNV backbone with the replacement of CHIKV E2 (ONNV/CHIKV E2), E1 (ONNV/CHIKV E1), or E2 and E1 (ONNV/CHIKV E2+E1). **b,** Titration of supernatant samples from THP-1 derived macrophages infected with CHIKV vaccine strain 181/clone 25, ONNV SG650, ONNV/CHIKV E2, ONNV/CHIKV E1, and ONNV/CHIKV E2+E1. Macrophages were inoculated with the viruses at MOI 5, and the supernatant samples were collected at 24 h.p.i for plaque assay analysis. Data are representative of 2 independent experiments. Mean values of biological duplicates measured in technical duplicates were plotted with SD. Asterisks indicate statistically significant differences as compared to ONNV (Two-way ANOVA and Dunnett’s multiple comparisons test: **, p≤0.01; ****, p≤0.0001). **c,** THP-1 derived macrophages were transfected with 0.5 μg RNA of CHIKV 181/clone 25, ONNV SG650, ONNV/CHIKV E2, ONNV/CHIKV E1, or ONNV/CHIKV E2+E1. Virion production was determined by intracellular (+) vRNA transcript levels and secreted infectious particle titers through RT-qPCR and plaque assay, respectively. Data are representative of 3 independent experiments. Mean values of biological duplicates measured in technical duplicates were plotted with SD. Asterisks indicate statistically significant differences as compared to ONNV (One-way ANOVA and Dunnett’s multiple comparisons test: **, p≤0.01; ****, p≤0.0001). **d,** Schematic representation of modified chimeras based on parental ONNV/CHIKV E1+E2 that contain hybrid E2 or E1. E2 has 3 domains: A, B connected to A and C by two flanking β-ribbon arches, and C. E1 has 3 domains: I, II, and III, with a fusion loop in II. Chimera containing hybrid E2 that has arch-B-arch-C (E2-I+E1), or only domain C (E2-II+E1) from ONNV. Chimera containing hybrid E1 has domains II and III (E2+E1-I), or only domain III (E2+E1-II) from ONNV. **e,** THP-1 derived macrophages were transfected with 0.5 μg RNA of ONNV SG650, ONNV/CHIKV E1 and chimeras (E2-I+E1, E2-II+E1), ONNV/CHIKV E2 and chimeras (E2+E1-I, E2+E1-II). Virion production was determined through RT-qPCR and plaque assays as described in Figure.3c. Data are representative of 4 independent experiments. Mean values of biological duplicates measured in technical duplicates were plotted with SD. Asterisks indicate statistically significant differences as compared to ONNV (One-way ANOVA and Dunnett’s multiple comparisons test: ***, p≤0.001; ****, p≤0.0001).

To map the viral determinants for virion production to specific domains, we strategically swapped in ONNV E2 or E1 domains in the context of ONNV/CHIKV E2+E1. Alphavirus E2 comprises three domains (A, B, and C) connected by β-ribbon arches, with domains A and B functioning in receptor binding and cellular attachment^31^. Alphavirus E1 consists of three β-barrel domains: I, II, and III, with the fusion peptide embedded in domain II critical for viral fusion and uncoating. We generated two ONNV/CHIKV E1 chimeras containing regions of CHIKV E2 (E2-I+E1, E2-II+E1) and two ONNV/CHIKV E2 chimeras containing regions of CHIKV E1 (E2+E1-I, E2+E1-II) (Figure.3d). We transfected macrophages with vRNAs of these chimeras in comparison with ONNV, ONNV/CHIKV E1, and ONNV/CHIKV E2 to measure virion production (Figure.3e). We found that only the chimeras containing CHIKV E2 without domain C or E1 without domain III restore virion production to significantly high levels. These results suggest that glycoprotein determinants crucial for virion production in macrophages may lie in CHIKV E2 domain B and flanking β-ribbon arches, and E1 domain II.

### Positively selected residues in E2 and E1 are essential for CHIKV production in macrophages

Recent SARS-CoV-2 studies have harnessed the power of complementary selection analyses to reveal residues under positive selection that might promote virus adaptation and expansion in human hosts^32–34^. CHIKV, like SARS-CoV-2, is a zoonotic virus well-adapted to humans. Therefore, we asked whether residues in the CHIKV glycoproteins have been under positive selection to overcome antiviral immunity and productively replicate in macrophages. We applied the same methodology from a highly cited SARS-CoV-2 study^32^. Combining the use of the fixed effects likelihood^35^ (FEL, p≤0.05) and mixed effects model of evolution^36^ (MEME, p≤0.05) methods, we analyzed 397 CHIKV structural polyprotein sequences from NCBI Virus^37^ database that were isolated from CHIKV-infected individuals globally (Figure.4a and Extended data.1a). FEL identified 4 amino acid residues in E2, 6K, and E1 under pervasive positive selection; MEME identified 14 residues in capsid, E2, 6K, and E1 under pervasive and episodic positive selection, including all 4 residues identified by FEL (Figure.4c and Extended data.1b).

**Figure 4.**
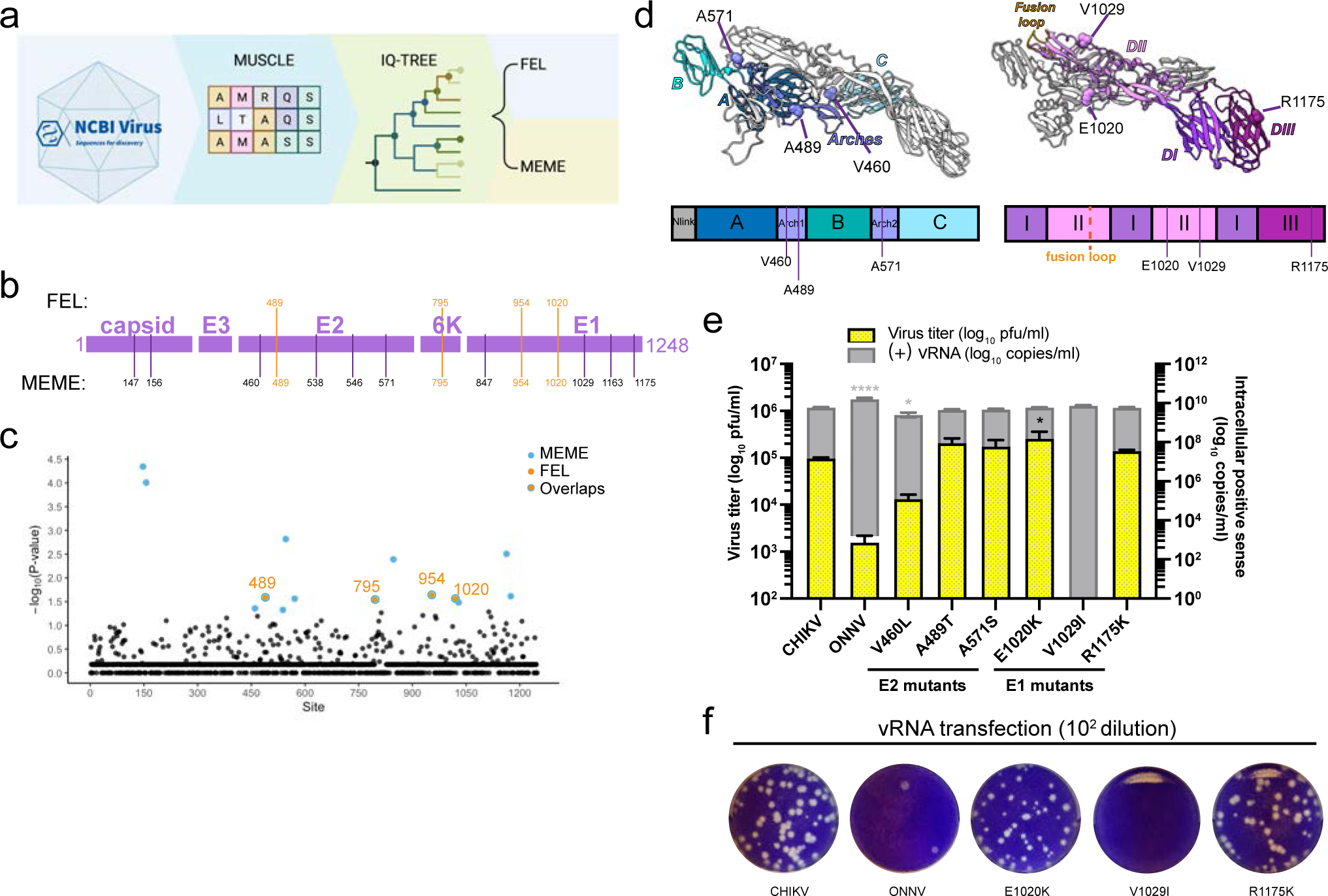
CHIKV E2 and E1 residues under positive selection are essential for virion production in human macrophages. **a,** The pipeline for analyzing natural selection in the evolution of CHIKV structural proteins in human hosts. 397 CHIKV sequences isolated from infected individuals globally were downloaded from the NCBI virus database, and structural polyprotein sequence alignment was performed by MUSCLE^82^. The phylogenetic tree of CHIKV was constructed based on maximum-likelihood (ML) optimality criterion with IQ-TREE^76, 83^. The sites under positive selection were identified using mixed effects model of evolution (MEME)^36^ and fixed effects likelihood (FEL)^35^. **b,** The positively selected sites identified by FEL or MEME are annotated in the structural polyprotein region of the CHIKV genome. Four of these sites (489, 795, 954, 1020) were identified with both methods and are colored in orange. **c,** The positively selected sites in CHIKV structural proteins detected by MEME or FEL are highlighted with their -log10 P-values determined by MEME. The statistically significant sites determined by MEME (p<0.05) are in blue, and the significant sites by both methods are in orange with blue circles. **d,** Visualization of CHIKV positively selected sites in the heterodimer formed by p62 (E3-E2) and E1. The structure of p62-E1 heterodimer was downloaded from PDB (3N40)^84^ and visualized in Chimera X^85^. Left: The positively selected sites V460, A489, and A571 are located in β-ribbon arches (purple) flanking domain B in E2. Right: The positively selected sites E1020 and V1029 are located in domain II (pink) in E1. The positively selected site R1175 is in domain III (magenta) in E1. **e,** Comparison of virion production of CHIKV positive selection site mutants in THP-1 derived macrophages. The positive selection site in E2 or E1 of CHIKV 181/clone 25 was mutated to the homologous residue in ONNV, respectively, to generate six CHIKV mutants (V460L, A489T, A571S, E1020K, V1029I, R1175K). Macrophages were transfected with 0.5 μg RNA of CHIKV, ONNV, or CHIKV positive selection site mutants, and virion production was determined by intracellular (+) vRNA transcript levels and secreted infectious particle titers as previously described. Data are representative of 3 independent experiments. Mean values of biological duplicates measured in technical duplicates were plotted with SD. Asterisks indicate statistically significant differences as compared to CHIKV (One-way ANOVA and Dunnett’s multiple comparisons test: *, p≤0.05; ****, p≤0.0001). **f,** Representative plaque images of CHIKV E1 positive selection site mutants (E1020K, V1029I, R1175K) in comparison with CHIKV and ONNV. Plaque assays were performed on supernatant samples from transfected THP-1 derived macrophages as mentioned in Figure 4e, and wells from 1:100 dilution are shown.

Interestingly, the positively selected sites identified by MEME were concentrated in E2 and E1 (Figure.4b and 4c). We found 3 residues in E2 (V460, A489, A571) and 3 in E1 (E1020, V1029, R1175) to be different between the CHIKV 181/clone 25 and ONNV SG650 strains (Extended data 1b). All the unique CHIKV residues are exposed on the surface of premature p62 (E3-E2)/E1 heterodimer before virion assembly (Figure.4d), suggesting potential host factor interaction sites.

To interrogate if these positively selected residues affect CHIKV production, we mutated them individually into the homologous residues in ONNV. We compared viral replication and production of these mutants (V460L, A489T, A571S, E1020K, V1029I, R1175K) with those of parental CHIKV in vRNA-transfected macrophages (Figure.4e). The V460L mutation decreases virus titers by about 1 log and significantly reduces intracellular (+) vRNA levels, which is likely due to unexpected cleavage of the E2 mutant (data not shown). Surprisingly, the V1029I mutation completely abrogates virion production in macrophages without affecting viral replication (Figure.4f), suggesting a defect in viral life cycle after genome replication. In contrast, both V460L and V1029I mutations attenuate viral replication and production in 293T and BHK-J cells (Extended data 2a and 2b). Interestingly, V460 and V1029 are in the E2 β-ribbon arch and E1 domain II, respectively, that were identified to be critical for virion production in Figure.3e. Taken together, the positively selected residue V1029 mediates efficient virion production in macrophages.

### Identification of cellular factors that interact with CHIKV glycoproteins in macrophages

Successful virion production requires the maturation of E2/E1 heterodimer for proper virion assembly which involves proteolytic processing of the precursor (E3-E2-6K-E1) to intermediate form (p62/E1), and finally to E2/E1 heterodimer in the secretory pathway^38–40^. To investigate intracellular macrophage factors that interact with the uncleaved precursors or mature glycoproteins to affect CHIKV production, we inserted a myc tag in CHIKV genome to label E2 N-terminally (CHIKV/myc-E2) that can also label the precursors in addition to E2/E1 heterodimers. We infected THP-1 derived macrophages in two independent experiments with either CHIKV/myc-E2 or untagged CHIKV 180/clone 25 (WT, negative control). We performed myc immunoprecipitation to enrich for myc-labeled precursor or mature glycoproteins, followed by MS analysis of the resultant protein mixtures to identify interactors (Figure.5a).

**Figure 5.**
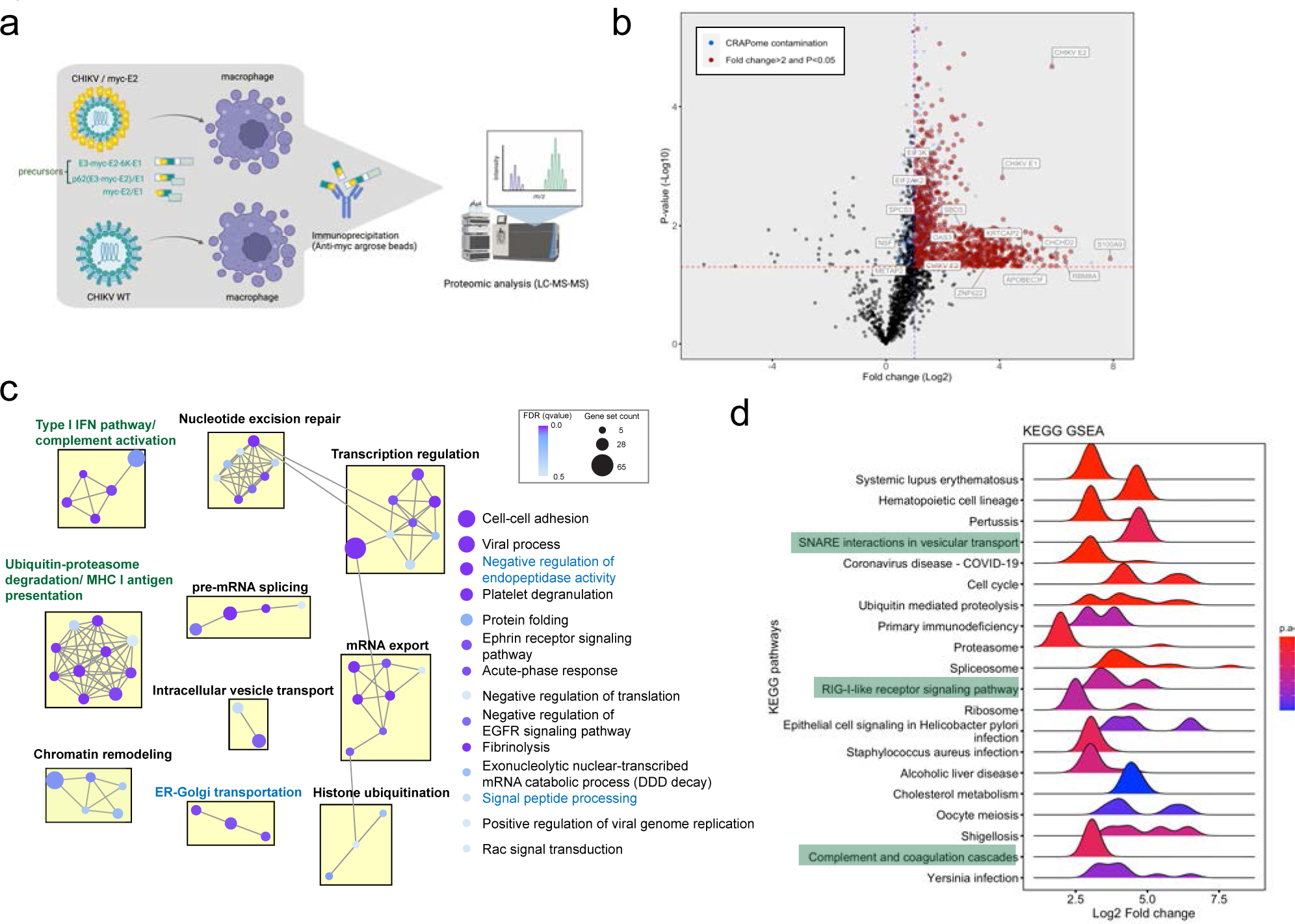
Identifying host factors interacting with CHIKV glycoproteins in infected human macrophages by affinity purification-mass spectrometry (AP-MS) **a,** The workflow of AP-MS analysis to identify host factors that interact with CHIKV glycoproteins. After 48 h infection with CHIKV/myc-E2, different forms of myc-tagged glycoproteins (E3-myc-E2-6K-E1, E3-myc-E2, myc-E2) were pulled down by anti-myc agarose beads from the infected cell lysates and submitted to LC-MS/MS analysis to identify co-immunoprecipitated host factors. The co-immunoprecipitated proteins from untagged CHIKV vaccine strain 181/clone 25 (CHIKV WT)-infected macrophages serve as negative control for proteomic analysis. **b,** Volcano plot depicting host interactors identified by mass spectrometry. The volcano plot is scattered by -log10 P-value (y-axis) and log2 expression fold change (FC) of proteins co-immunoprecipitated from CHIKV/myc-E2 infected cells with respect to the proteins from CHIKV WT infected cells (x-axis). The dashed cut-offs of adjusted P-value and expression fold change are 0.05 (-log10P-value=1.30103) and >2 (log2FC=1), respectively. CHIKV glycoproteins (E3, E2, E1) and host factors for further investigation in Figure 6a are annotated in box. **c,** Enrichment map that summarizes over-represented biological processes of identified host factors in groups. The enriched proteins identified by mass spectrometry were clustered by biological processes and organized into a network with edges connecting overlapping gene sets to reveal the functional module. **d,** Gene Set Enrichment Analysis (GSEA) of top 20 KEGG pathways in identified host factors summarized in ridge plot. All the identified host factors are ranked according to the log2 expression fold change of proteins co-immunoprecipitated from CHIKV/myc-E2 infected macrophages with respect to proteins from CHIKV 181/clone 25 infected macrophages (x-axis). The significance of the KEGG enrichment is shown in continuous color scale based on the adjusted p values (FDR). The histogram in each KEGG term is defined by the number of genes with a specific log2 fold change value.

**Figure 6.**
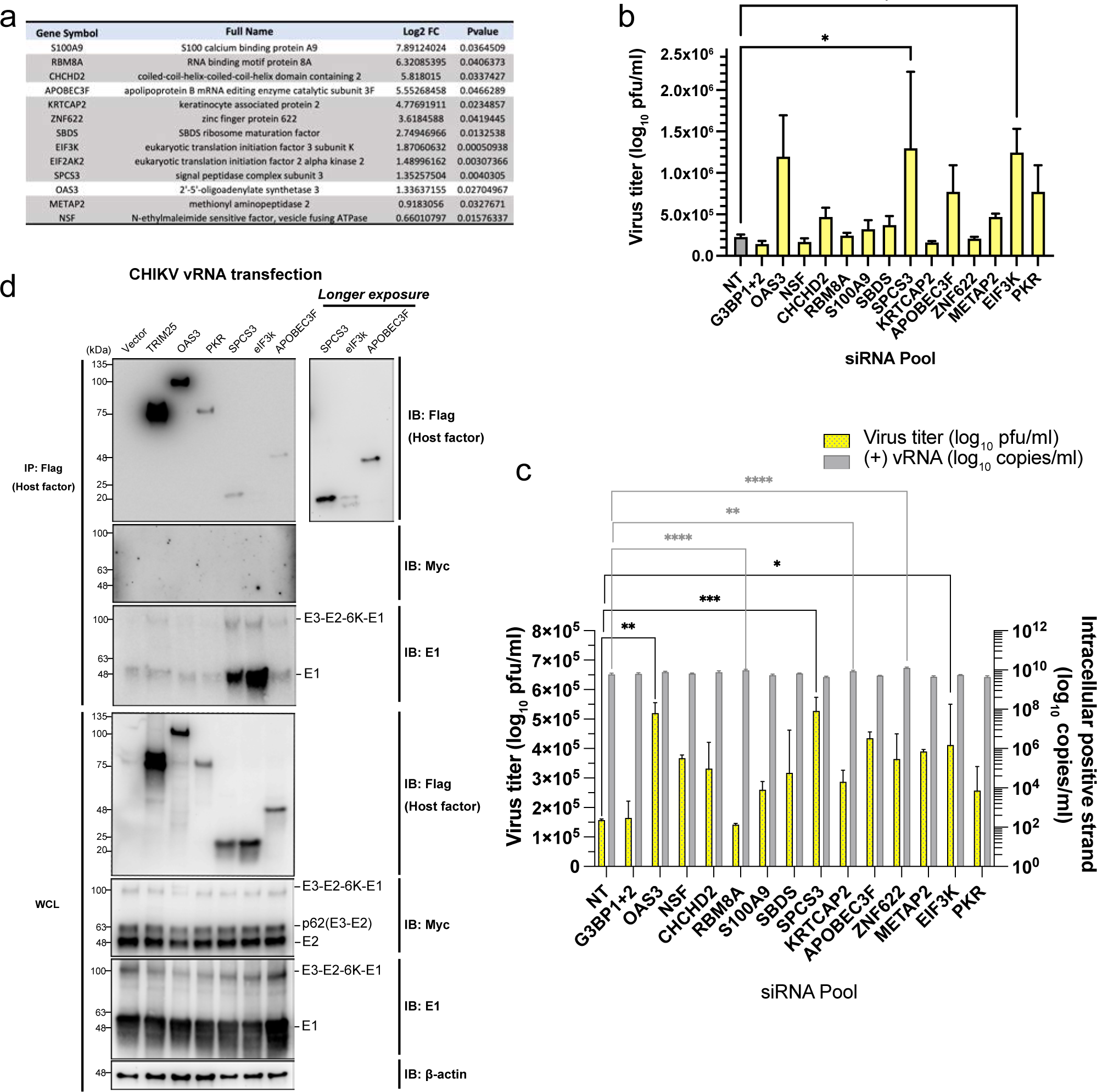
CHIKV E1 interacts with host factors that block virion production in human macrophages. **a,** Table of identified host factors that were chosen for siRNA knockdown assays in Figure. 6b and 6c. The gray-highlighted genes are significantly detected in 2 independent AP-MS experiments. **b-c,** Evaluation of CHIKV infection **(b)** and production **(c)** in human macrophages with OAS3, NSF, CHCHD2, RBM8A, S100A9, SBDS, SPCS3, KRTCAP2, APOBEC3F, ZNF622, METAP2, EIF3K, or PKR knocked down. THP-1 derived macrophages were transfected with pooled siRNAs targeting specific host factors or nontargeting siRNAs (NT) for 48 h. The cells were then infected with CHIKV (MOI 5) **(b)** or transfected with CHIKV vRNA **(c)** for 24 h. The supernatant virus titers from cells treated with siRNAs targeting host factors were determined by plaque assay and compared to the titers from cells treated with NT siRNA to assess the anti- or pro-viral effects of specific host genes on CHIKV production. G3BP1 and G3BP2 (G3BP1+2) known to be proviral for CHIKV replication were knocked down together as control. Data are representative of 2 independent experiments. The mean values of biological duplicates measured in technical duplicates were plotted with SD (One-way ANOVA and Dunnett’s multiple comparisons test: *, p≤0.05; **, p≤0.01; ***, p≤0.001, ****, p≤0.0001). **d,** 293T cells were transfected with plasmids expressing 3xFLAG-tagged host factors (TRIM25, OAS3, PKR, SPCS3, eIF3k, and APOBEC3F) or empty vector control for 24 h and later transfected with vRNA of CHIKV/myc-E2. The cells were lysed and immunoprecipitated by anti-FLAG agarose beads. Immunoblot was probed to check for E2/E1 binding to these host factors. TRIM25-3xFLAG was transfected into 293T cells for immunoprecipitation control. Data are representative of 3 independent experiments.

We identified 1157 proteins (Log2FC>0, p<0.05) in the second experiment to be significantly enriched in CHIKV/myc-E2-infected cells compared to CHIKV WT infected cells (Figure.5b and Supplementary information 3). In addition to the bait protein E2 (Log2FC=5.84, p=2.11E-5), we also detected E1 (Log2FC=4.09, p=1.59E-3) and E3 (Log2FC=2.99, p=3.08E-2) as expected. Gene ontology enrichment analysis visualized with enrichment map (Figure.5c) showed that host factors are highly enriched in grouped or singleton biological processes (BPs) relevant to the secretory pathway in blue (negative regulation of endopeptidase activity, signal peptide processing, ER-Golgi transportation, intracellular vesicle transport) and innate immune responses in green (type I IFN pathway/complement activation, antigen presentation) (Figure.5c). We also performed KEGG analysis^41–43^ highlighting that CHIKV glycoproteins manipulate secretory pathway and innate immune responses in green, which is consistent with BP analysis (Figure.5d). We then deciphered prominent biological processes through STRING PPI network^44^ and discovered protein complexes related to signal peptidases and type I IFN pathway (Extended data 3). Interestingly, signal peptidases are hijacked for polyprotein maturations of several viruses including alphavirus^45, 46^, however, the peptidases involved are still unknown.

### CHIKV E1 binding proteins exhibit potent anti-CHIKV activities

We next inquired whether the host factors interacting with the CHIKV glycoproteins are proviral or antiviral. We selected 13 host factors for further investigation, including 10 hits identified in both AP-MS experiments, classical ISGs (APOBEC3F, OAS3), and a myeloid-specific gene (S100A9) which is an endogenous ligand for toll-like receptor 4 (TLR4)^47, 48^ (Figure.6a). We knocked down these genes with pooled siRNAs (Extended data 4a) in macrophages, followed by CHIKV infection (Figure.6b). We included non-targeting (NT) siRNA as negative control and siRNAs targeting pro-CHIKV factors G3BP1 and G3BP2^49, 50^ as positive control. Knockdown of most of the host factors led to elevated CHIKV titers in macrophages compared to NT-transfected cells, except for G3BP1+2 knockdown, indicating that many of the candidate E2 interactors have antiviral activities. In addition to the previously reported anti-CHIKV restriction factors OAS3 and PKR^51–53^, knockdown of the host genes SPCS3 and EIF3K significantly restores virion production by about 5-fold. To confirm that the antiviral activities observed in Figure.6b are specific to a step after viral entry, we knocked down the same host factors in macrophages followed by transfection of CHIKV vRNA (Figure.6c). We found that silencing of most of the genes enhances virion production in vRNA-transfected macrophages. CHIKV production in macrophages with OAS3, SPCS3, and EIF3K knockdown is significantly higher than that in NT-transfected cells, despite similar intracellular vRNA levels.

To confirm the interaction of CHIKV glycoproteins with host proteins demonstrating antiviral activities (OAS3, SPCS3, eIF3k, APOBEC3F, and PKR, Figure.6b-c), we transfected 293T cells with plasmids expressing 3XFLAG-tagged host factors, followed by transfection with CHIKV vRNA (Figure.6d) or CHIKV polyprotein (E3-myc-E2-6K-E1) expressing plasmid (Extended data 4b). The host factors were pulled down to probe for glycoproteins in precursor or mature forms. We consistently detected strong binding of E1 and moderate binding of E3-E2-6K-E1 to SPCS3 and eIF3k, which suggests that host factors interact with precursors and E2/E1 heterodimer through binding to E1. Meanwhile, we did not observe any binding of E2 or p62 to these selected host factors. Taken together, these results demonstrate that E1 interferes with intrinsic host blockades through sequestering unconventional antiviral proteins.

## Discussion

Macrophages are important cellular reservoirs for persistent CHIKV infection; however, the underlying mechanisms are largely unexplored. In this study, we interrogated the CHIKV determinants that hijack macrophages as virion disseminators. We first demonstrated that both CHIKV glycoproteins E2 and E1 mediate efficient virion production from infected macrophages through comparative infection with CHIKV-ONNV chimeras. By performing selection analysis on sequences of human CHIKV isolates from NCBI Virus^37^, we identified E2-V460 and E1-V1029 to be associated with elevated CHIKV production. We then uncovered new antiviral factors SPCS3 and eIF3k by AP-MS analysis that bind strongly to E1 and restrict CHIKV production in macrophages. Our results suggest that positive selection of E2/E1 potentially driven by viral glycoprotein antagonism of cellular restrictions contributes to efficient CHIKV production in macrophages.

The six positive selection sites (460, 489, 571, 1020, 1029, 1175) in the structural proteins that are divergent between CHIKV (vaccine strain 181/clone 25) and ONNV (strain SG650) are primarily in the E2 β-ribbon arches and E1 domain II, away from the embedded fusion peptide. All of the differential selection sites are exposed on the outer surface of p62-E1 that is accessible for interactions with host factors in the secretory pathway and can influence E2/E1 heterodimer maturation. Our results showed that mutating CHIKV E2-V460 and E1-V1029 into ONNV residues disrupts virion production with the former causing unexpected cleavage of E2, suggesting that the positively selected valine residues in these sites might counteract host restriction factors targeting maturation. Consistent with that, silencing of E2/E1 interactors identified by AP-MS revealed primarily antiviral activities at a late step in the viral life cycle after genome replication.

Components of the adaptive immune response, such as inhibitory antibodies and T cells, can also select for escape mutations in viral glycoproteins^25^. The epitopes of currently characterized human CHIKV neutralizing antibodies or broadly anti-alphavirus antibodies are mainly mapped to E2 domains A and B, responsible for receptor binding and cellular attachment, and E1 domain II, proximal to or within the fusion loop^54–56^. None of these reported antibodies target the six differential selection sites in CHIKV 181/clone 25 (Figure.4d), suggesting that these residues are likely selected by host restriction factors.

Among the candidate CHIKV glycoprotein interactors we identified, SPCS3 and eIF3k have inhibitory activities against CHIKV production in human macrophages. SPCS3 is one of the core components of the endoplasmic reticulum-associated signal peptidase complex (SPC)^57^, which cleaves signal peptides during the translocation of protein precursors in the ER^58, 59^. The signal peptidases are presumably usurped by flaviviruses, bunyaviruses, and alphaviruses for poly-glycoprotein cleavage^45, 46, 60^. However, it is unknown what exact peptidase releases p62, 6K, and E1 from the alphavirus poly-glycoprotein precursor^61^. A previous genome-wide CRISPR knockout screen uncovered both SPCS1 and SPCS3 as proviral factors for flavivirus infection, and depletion of SPCS1 led to inefficient polyprotein cleavage disrupting flavivirus production^60^. Surprisingly, we found that SPCS3 exhibits anti-CHIKV activity and strongly associates with CHIKV E1. SPCS3 overexpression does not affect CHIKV poly-glycoprotein cleavage (Figure.6d), suggesting novel peptidase-independent antiviral activities. For the first time, we demonstrated functional dualities of SPC proteins in different virus infection systems.

eIF3k is a component of the eIF3 complex that binds 40S subunit, and is involved in almost all steps of translation initiation^62, 63^. Through protein cluster analysis of our AP-MS results (Extended Data 3), we also detected host factors associated with eIF3k, including eIF2a, a crucial subunit of eIF2, and receptor for activated C kinase 1 (RACK1). eIF2a phosphorylation can be induced by double stranded RNA formed during viral replication, leading to translation shutoff that suppresses viral gene expression^64^. Therefore, it is plausible that eIF3k antiviral activity is related to eIF2a-mediated translation inhibition. Moreover, previous studies indicated that RACK1 acts as a kinase scaffold to potentially regulate the phosphorylation status of several eIF3 subunits^65, 66^. Future studies are needed to determine whether eIF3k regulates viral translation in a RACK1-dependent manner.

In summary, our study has uncovered CHIKV determinants for virion production in macrophages in the light of evolutionary selection and proposes a novel role for viral glycoproteins as antagonists of host antiviral immunity. Overall, this research provides promising viral and cellular targets for therapeutic intervention to strengthen the antiviral status of macrophages and eliminate CHIKV reservoirs.

## Methods

### Cell culture, viruses, and infections

BHK-21 cells (American Type Culture Collection (ATCC)) were maintained in Minimum Essential Media (MEM, Gibco) supplemented with 7.5% fetal bovine serum (FBS, VWR). HEK-293T cells (ATCC) were maintained in Dulbecco’s Modified Eagle Medium (DMEM, VWR) supplemented with 10% FBS. THP-1 human monocytes (ATCC) were maintained in Roswell Park Memorial Institute 1640 Medium (RPMI 1640, Gibco) supplemented with 10% FBS, 1X penicillin/streptomycin (P/S, Fisher Scientific), 1X non-essential amino acids (NEAA, Fisher Scientific), and 0.05 mM β-mercaptoethanol (Sigma-Aldrich).

The infectious clone plasmids of enhanced GFP (EGFP)-expressing or unlabeled CHIKV (vaccine strain 181/clone 25), EGFP-expressing or unlabeled ONNV (strain SG650), EGFP-expressing SINV (TE/5’2J/GFP), and EGFP-expressing RRV (strain T48) have been previously reported^67–71^ Propagations and titrations of virus stocks were generated in BHK-21 cells as previously described^72, 73^. To infect THP-1 derived macrophages or primary monocyte derived macrophages, viruses were diluted in DPBS supplemented with 1% human AB serum (Omega Scientific) and 1% P/S, and added to cells at a multiplicity of infection (MOI) of 5 plaque-forming units (PFU)/cell. Typically, infection was carried out in a 12-well or 24-well plate with 5 × 10^5^ or 2.5 × 10^5^ macrophages seeded per well. Cells were incubated with virus for one hour and washed twice with PBS to remove the virus. Freshly made media was then added to cells, and supernatant samples were collected at the indicated timepoints for plaque assay as previously described.

### Monocyte differentiation and transfection

THP-1 human monocytes were differentiated into macrophages through a 24-hour (h) stimulation with 50 ng/mL phorbol 12-myristate 13-acetate (PMA, Sigma-Aldrich) in RPMI 1640 supplemented with 10% human AB serum, 1X NEAA, 1X P/S followed by a 24 h rest in human-serum containing RPMI 1640.

Human primary peripheral blood mononuclear cells (PBMCs) were obtained from donors through the UCLA/CFAR Virology Core Lab (grant number 5P30 AI028697). The RosetteSep™ Human Monocyte Enrichment Cocktail (STEMCELL Technologies) was used to purify monocytes from the PBMCs. To differentiate the purified monocytes to macrophages, the monocytes were cultured in ImmunoCult™-SF Macrophage Medium (STEMCELL Technologies) supplemented with 50 ng/uL Human Recombinant M-CSF (STEMCELL Technologies) for 4 days. After differentiation, the macrophages were infected as described in the previous section.

### siRNA and viral RNA transfection

For gene silencing, three unique Ambion Silencer siRNAs (Thermo Fisher Scientific) targeting 13 host factors identified by AP-MS were pooled and transfected into THP-1 macrophages at a final concentration of 25 nM. To simultaneously knock down G3BP1 and G3BP2 as positive control, two unique Ambion Silencer siRNAs respectively targeting G3BP1 and G3BP2 were pooled (25 nM) and transfected into THP-1 macrophage. The same amount of non-targeting siRNA (Thermo Fisher Scientific) was transfected into THP-1 macrophages as negative control. siRNA transfections were performed with TransIT-X2 Transfection Kit (Mirus Bio) following manufacturer’s instructions. Downstream assays were conducted 48 h after transfection.

To observe viral production in transfected macrophages, 500 ng of viral genomic RNA was transfected per well in 12-well plates through the TransIT®-mRNA Transfection Kit (Mirus Bio) following manufacturer’s instructions.

### The construction of CHIKV-ONNV chimeras and point mutants

All the primers and restriction sites used in chimeras, mutants, and reporter virus constructions mentioned below are listed in Tables 1, 2, and 3 in **Supplementary information 2** respectively.

To construct Chimera I, gene regions amplified from the parental CHIKV vaccine strain 181/clone 25 and ONNV SG650 strains were fused into two chimeric fragments, Fragment 1 and Fragment 2, through PCR overlap extension (**Supplementary information 1**). Fragment 1 was inserted into the CHIKV 181/clone 25 backbone to generate an intermediate chimera with parts of nsP4 and capsid from ONNV. The fragment from ONNV subgenomic promoter to the end of CHIKV poly(A) tail was digested from the intermediate chimera and inserted into ONNV backbone with Fragment 2 to obtain Chimera I.

To generate Chimera III, we first used overlapping PCR to generate Fragment 3 to replace the equivalent region in CHIKV 181/ clone 25 to obtain the CHIKV/ONNV 5’UTR backbone. We then used the NEBuilder HiFi DNA Assembly Kit to ligate the CHIKV/ONNV 5’UTR backbone with CHIKV subgenomic promoter and capsid (Fragment 4) and ONNV E3 to the end of the poly(A) tail (Fragment 5) (**Supplementary information 1**). Both Fragments 4 and 5 contained overlapping overhangs for HiFi ligation.

The cloning of Chimera II was based on Chimera I. We amplified the region from the CHIKV subgenomic promoter to the PspXI site in E2 with overlapping overhangs and used the NEBuilder HiFi DNA Assembly Kit to ligate the amplified product to the digested Chimera I backbone. To generate Chimera IV, we amplified the region from the ONNV subgenomic promoter to the intrinsic BamHI site in ONNV E2. We then used T4 ligase (New England Biolabs) to ligate the amplified fragment with digested Chimera III backbone.

The other chimera clone plasmids (Chimera III-I, III-II, III-III, ONNV/CHIKV E1, ONNV/CHIKV E2, ONNV/CHIKV E1+E2, E2-I+E1, E2-II+E1, E2+E1-I and E2+E1-II) were generated in a similar fashion through multiple fragment ligations with the NEBuilder HiFi DNA Assembly Kit.

To construct the CHIKV positive selection site mutants (V460L, A489T, A571S, E1020K, V1029I, and R1175K), the region containing E2 or E1 was amplified from CHIKV 181/clone 25 and inserted into pCR-Blunt II-TOPO vector (Thermo Fisher Scientific) according to manufacturer’s instructions. Corresponding site-directed mutagenesis was conducted on the intermediate TOPO constructs with specific mutation primers by using the Q5® Site-Directed Mutagenesis Kit (New England BioLabs). The mutated E2- or E1-containing fragments were digested from the TOPO constructs through intrinsic viral restriction sites and inserted back into CHIKV through T4 ligation.

To construct CHIKV with myc-tagged E2 (CHIKV/myc-E2), the myc tag was inserted between E3 and E2 through the NEBuilder HiFi DNA Assembly Kit. Fragment 6 was amplified from parental CHIKV 181/clone 25, containing the region from the subgenomic promoter in nsp4 to the end of E3. The segment of E2, from the start of E2 to the second NdeI site, was amplified from CHIKV 181/clone 25 as Fragment 7. The reverse primer of Fragment 6 and forward primer of Fragment 7 incorporates the myc tag into CHIKV 181/clone 25 through three-fragment assembly (**Supplementary information 1**).

### Construction of host factor and CHIKV structural polyprotein plasmids

All the primers and restriction sites used in the construction of host factor and CHIKV structural polyprotein plasmids are listed in Table 2 in **Supplementary information 1**, respectively.

The cellular mRNA from THP-1 cells was reverse transcribed with oligo-dT primer through the Protoscript II First Strand cDNA Synthesis Kit (New England Biolabs) after TRIzol (Thermo Fisher Scientific) extraction. The host genes OAS3, PKR, SPCS3, EIF3K, and APOBEC3F were amplified with specific primers containing regions overlapping the pcDNA3.1-3xFLAG vector. The cDNAs of host factors were then incorporated into the pcDNA3.1-3xFLAG vector digested with NotI and XbaI sites through the NEBuilder HiFi DNA Assembly Kit.

To construct the plasmid for CHIKV structural glycoprotein following capsid cleavage (pcDNA3.1-E3-myc-E2-6K-E1), the sequence spanning the beginning of E3 to the end of E1 was amplified from CHIKV/myc-E2 with primers containing overlapping regions with the pcDNA3.1 vector and incorporated into pcDNA3.1 through NEBuilder HiFi DNA Assembly Kit.

### Quantitative reverse transcription PCR (RT-qPCR)

Cells were lysed with TRIzol reagent (Thermo Fisher Scientific) followed by extraction of total RNAs through the Direct-zol RNA Microprep Kit (Zymo Research) according to manufacturer’s instructions. To enhance assay specificity, tagged reverse transcription primers targeting viral genes were used to synthesize viral cDNAs from total RNAs. The transcribed cDNAs were then quantified by SYBR Green or TaqMan qPCR.

The SYBR Green assay was used to evaluate the copy number of (+) vRNA in the samples. To generate standard curve transcripts, full-length CHIKV E1 and partial ONNV E1 (SG650 bp 10092-11361) sequences were amplified with reverse primers containing the SP6 promoter and inserted into the pcDNA3.1 vector with an inherent T7 promoter at the 5’ terminal end. The (+) and (-) standard curve transcripts were synthesized with T7 polymerase using HindIII-linearized plasmid and Sp6 polymerase using NheI-linearized plasmid, respectively, through the MAXIscript™ SP6/T7 Transcription Kit (Thermo Fisher Scientific). The cDNAs of (+) standard curve transcripts and viral RNA in the samples were reverse transcribed with a reverse E1 primer containing a nongenomic tag sequence^74^ 5’-CAGACAGCACTCGTTCGTACAC-3’ through the Protoscript II First Strand cDNA Synthesis Kit (New England Biolabs). The (+) standard curve cDNAs were then serially diluted ten-fold from 10^-1^ to 10^-8^ and run through the SYBR Green assay (New England Biolabs) together with sample cDNAs. Specific forward primer targeting E1 and a reverse primer targeting the nongenomic tag were used in 20 ul SYBR Green reaction with 1x Luna qPCR Dye (New England Biolabs) according to manufacturer’s instructions. The reactions were run under the cycling conditions as previously reported^73^.

The TaqMan assay was performed to determine the copy numbers of (-) vRNAs in the samples. The standard curve of (-) strand nsP1 from CHIKV or ONNV, tagged reverse transcription primers, qPCR primers, and TaqMan probes were designed and generated as previously described^75^. Briefly, after in vitro transcription of (-) nsP1 transcripts from standard curve plasmids, the cDNAs of (-) nsP1 transcripts and viral RNA in the samples were synthesized with a forward nsP1 primer containing a unique tag sequence 5’-GGCAGTATCGTGAATTCGATGC-3’ by the Protoscript II First Strand cDNA Synthesis Kit. The appropriate reverse nsP1 primer, tag-specific forward primer, and FAM-labeled TaqMan probe were used in viral negative strand quantification with Luna Universal Probe qPCR Master Mix (New England Biolabs). The reactions were run under the cycling conditions as follows: initial denaturation step at 95°C for 1 min followed by 40 cycles of 95°C for 15 s and 60°C for 30 s. Data collection occurs during the 60°C extension step.

Both SYBR Green and TaqMan reactions were performed in technical duplicates of cDNA samples from biological duplicate wells. All qPCR reactions were run on the CFX96 OPUS (Bio-Rad). The total copy number of viral RNA was determined by using the standard curve method. All the primers used in qPCR assays are listed in Table 4 in supplementary information.

### Positive selection analysis

Chikungunya virus (taxid: 37124) structural polyprotein sequences were downloaded from the NCBI Virus database. Sequences that were not isolated from a human host, less than 10,000 nucleotides in length, or had more than 0.5% of ambiguous characters were excluded; 556 sequences remained.

To guide the nucleotide alignment, the sequences were first translated to amino acids with HyPhy’s Codon-aware MSA program (pre-msa). The amino acids were aligned with MUSCLE^1^ and used to align the nucleotide sequences with HyPhy’s Codon-aware MSA program (post-msa). A maximum likelihood phylogenetic tree was constructed by IQ-TREE^76^. By using HyPhy’s FEL^35^ and MEME^36^ methods, positive selection analyses were performed on 397 sequences after exclusion of duplicates from the original 556 sequences.

### Co-immunoprecipitation and immunoblot

To prepare samples for AP-MS, THP-1 monocytes were differentiated into macrophages in 36 15-cm dishes with 2 × 10^7^ cells per dish. Half of the dishes were either infected with CHIKV vaccine strain 181/clone 25 or CHIKV/myc-E2 at an MOI of 5 PFU/cell. Forty-eight hours later, cells in each dish were lysed with 2 mL NP40 lysis buffer (100 mM Tris-HCl (pH 8.0), 5 mM EDTA, 150 mM NaCl, 0.1% NP-40, 5% glycerol) supplemented with 1X PMSF, 2X PPI, 1 uM DDT, and Complete EDTA-free protease inhibitor mixture tablet (Roche). Cell lysates from every 6 dishes under same treatment were combined and further centrifuged at 14000*xg* for 15 mins. The clarified supernatants were incubated with anti-myc agarose beads (EZview™ Red Anti-c-Myc Affinity Gel, Millipore) for 4 hours at 4°C. After washing with NP40 lysis buffer 4 times, proteins were eluted with urea buffer (8M urea, 100 mM Tris HCl (pH 8)) for mass spectrometry analysis.

To validate our mass spectrometry hits, HEK-293T cells were seeded in 6-well plates at a starting density of 1.5 × 10^5^ cells/well, followed by transient transfection with plasmids expressing 3xFLAG-tagged host factors (OAS3, PKR, SPCS3, EIF3K, and APOBEC3F), empty vector, or control plasmid expressing 3xFLAG-tagged TRIM25 through X-tremeGENE9 (Millipore Sigma). Twenty-four hours later, the cells were either transfected with CHIKV/myc-E2 viral RNA (2 μg/ well) through TransIT®-mRNA Kit or CHIKV structural polyprotein plasmid (1 μg/ well) through X-tremeGENE9. Cells in each well were lysed with 300 μl NP40 lysis buffer 24 hr later and clarified by centrifugation as mentioned above. Immunoprecipitation of FLAG-tagged host factors in clarified supernatants with anti-FLAG agarose beads (EZview™ Red Anti-FLAG M2 Affinity Gel, Sigma-Aldrich) was performed at 4°C for 45 minutes. After 4x washing, proteins were directly eluted with Laemmli Sample Buffer (Bio-Rad) containing 5% 2-mercaptoethanol and denatured by boiling.

Proteins were resolved by SDS-PAGE in 4-15% precast Mini-PROTEAN TGX Gels (Bio-Rad) in conventional Tris/Glycine/SDS buffer. Proteins were blotted to PVDF membrane (Bio-Rad) and detected with primary antibodies and HRP-conjugated secondary antibodies listed in Table 5 in **Supplementary information 2**. Immunoblots were imaged by chemiluminescence with the ProSignal Pico ECL Reagents (Genesee Scientific) on a ChemiDoc (Bio-Rad).

### Mass Spectrometry and data analysis

Two independent AP-MS experiments were performed to identify macrophage proteins that interact with CHIKV glycoproteins. For mass spectrometry, protein disulfide bonds were subjected to reduction using 5 mM Tris (2-carboxyethyl) phosphine for 30 min and free cysteine residues were alkylated by 10 mM iodoacetamide for another 30 min. Samples were diluted with 100 mM Tris-HCl at pH 8 to reach a urea concentration of less than 2 M, and then digested sequentially with Lys-C and trypsin at a 1:100 protease-to-peptide ratio for 4 and 12 hours, respectively. The digestion reaction was terminated by the addition of formic acid to 5% (vol/vol) with centrifugation. Finally, samples were desalted using C18 tips (Thermo Scientific, 87784) and dried in a SpeedVac vacuum concentrator, and reconstituted in 5% formic acid for LC-MS/MS processing.

Tryptic peptide mixtures were loaded onto a 25 cm long, 75 μm inner diameter fused-silica capillary, packed in-house with bulk 1.9 μM ReproSil-Pur beads with 120 Å pores as described previously^77^. Peptides were analyzed using a 140 min water-acetonitrile gradient delivered by a Dionex Ultimate 3000 UHPLC (Thermo Fisher Scientific) operated initially at 400 nL/min flow rate with 1% buffer B (acetonitrile solution with 3% DMSO and 0.1% formic acid) and 99% buffer A (water solution with 3% DMSO and 0.1% formic acid). Buffer B was increased to 6% over 5 min at which time the flow rate was reduced to 200 nl/min. A linear gradient from 6-28% B was applied to the column over the course of 123 min. The linear gradient of buffer B was then further increased to 28-35% for 8 min followed by a rapid ramp-up to 85% for column washing. Eluted peptides were ionized via a Nimbus electrospray ionization source (Phoenix S&T) by application of a distal voltage of 2.2 kV.

All label-free mass spectrometry data were collected using data dependent acquisition on Orbitrap Fusion Lumos Tribrid mass spectrometer (Thermo Fisher Scientific) with an MS1 resolution of 120,000 followed by sequential MS2 scans at a resolution of 15,000. Data generated by LC-MS/MS were searched using the Andromeda search engine integrated into the MaxQuant 2 bioinformatic pipelines against the Uniprot Homo sapiens reference proteome (UP000005640 9606) and then filtered using a “decoy” database-estimated false discovery rate (FDR) < 1%. Label-free quantification (LFQ) was carried out by integrating the total extracted ion chromatogram (XIC) of peptide precursor ions from the MS1 scan. These LFQ intensity values were used for protein quantification across samples. Statistical analysis of differentially expressed proteins was done using Bioconductor package ArtMS3. Samples were normalized by median intensity.

Due to higher protein abundance, results from our second AP-MS experiment were visualized by volcano plot (ggplot2.tidyverse.org) to show host factors that were significantly enriched by myc pulldown in CHIKV/myc-E2 infected macrophages. To perform gene ontology analysis, candidate host interactors were first filtered by the cut-offs of p value < 0.05 and Log_2_ fold change > 0, based on the comparison of CHIKV/myc-E2 treatment group to CHIKV 181/clone 25 treatment group. We then used the CRAPome (Contaminant Repository for Affinity Purification) database (crapome.org) to remove potential contaminant proteins by a cutoff of ≥ 200 appearances in 716 recorded experiments. The filtered host factors were submitted to The Database for Annotation, Visualization and Integrated Discovery (DAVID: david.ncifcrf.gov) to analyze the enriched biological process (BP) categories. To have an intuitive view of all the BP categories, EnrichmentMap^78^ in Cytoscape^79^ was used to generate the network of all the BP enrichment results. The KEGG pathway analysis on host factors was performed by the latest online KEGG database (kegg.jp) downloaded in ClusterProfiler^42^, and the distribution of core enriched host factors for KEGG categories were visualized through ridgeplot in ClusterProfiler. To detect protein complexes in protein-protein interaction network with AP-MS results, filtered host proteins were reconstructed in STRING network^44^ with confidence score 0.7 and sequentially clustered by MCODE^80^ through clusterMaker^81^ to classify representative protein complexes.

### Flow cytometry

After 24 hr incubation with EGFP-labeled alphaviruses, primary human monocyte derived macrophages were detached from 12-well plate by using accutase (Stemcell Technologies). Digested macrophages were washed with PBS for 2 times in 96-well plates and fixed in fixation buffer (1% paraformaldehyde (PFA), 1% FBS in PBS). The intracellular EGFP expressions were detected by MACSQuant Analyzer (Miltenyi Biotec) with minimum collection of 20,000 events per sample. The results were analyzed through FlowJo (Tree Star).

## Acknowledgements

We especially thank Dr. Stephen Higgs (Kansas State University) for help with chimeric virus construction and Dr. Mehdi Bouhaddou (University of California, Los Angeles (UCLA)) for suggestions on proteomic analysis. We express our gratitude to Dr. Sergei L.Kosakovsky Pond (Temple University), his students Jordan Zehr and Alexander Lucaci for their insights on positive selection analysis. We thank Dr. Joyce Jose and her student Zeinab Elmasri from Pennsylvania State University as well as Dr. Graham Simmons and Dr. Jing Jin from Vitalant Research Institute for help with the construction of CHIKV E2 reporter virus. We also thank Dr. Oliver Fregoso (UCLA) for his comments on this study. We thank the UCLA Proteome Research Center for their services. We thank the qPCR and flow cytometry platforms provided by UCLA AIDS Institute which is supported by the James B. Pendleton Charitable Trust and the McCarthy Family Foundation. We also thank UCLA/CFAR Virology Core Lab (grant number 5P30 AI028697) for providing human primary monocytes. This work was supported in part by NIH R01AI158704 (MMHL), UC Cancer Research Coordinating Committee Faculty Seed Grant (CRN-20-637544; MMHL), and Sydney Finegold Post-Doctoral Fellow Award (ZY). JAW was supported by NIH GM089778.

## Supplementary information

Supplementary information 1: Schematics for ONNV-CHIKV chimera and CHIKV/myc-E2 construction.

Supplementary information 2: Excel sheets of primers, vectors, restriction sites and antibodies. Supplementary information 3: AP-MS protein list.

**Extended data 1.**
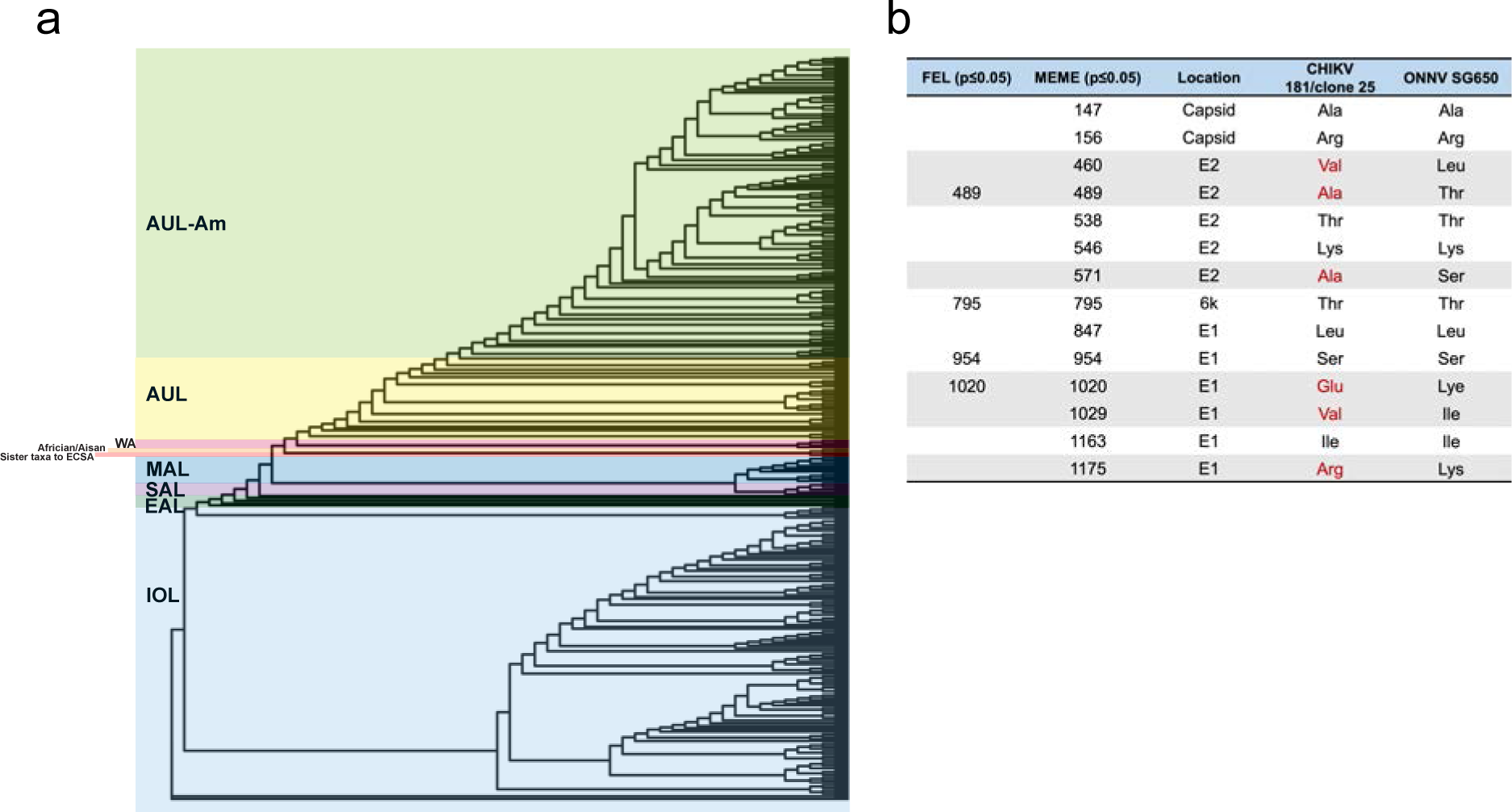
**a,** Phylogenetic tree constructed by IQ-tree^76^ using an alignment of the CHIKV structural polyprotein. The tree was visualized by ggtree^86^. Tree branches were colored according to the latest CHIKV lineage classification^87^ used in CHIKVnext v3 (nextstrain.org/groups/ViennaRNA/CHIKVnext/v3.0). AUL-Am: Asian Urban +American Lineage; AUL: Asian Urban Lineage; EAL: Eastern African Lineage; IOL: Indian Ocean Lineage; MAL: Middle African Lineage; SAL: South American Lineage; WA: Western African Lineage. **b,** Table of evolutionary positive selection sites in CHIKV structural polyprotein that were identified by FEL and MEME. The amino acid positions in structural polyprotein and CHIKV genes were annotated. The positively selected CHIKV amino acids that are different from the homologous residues in ONNV were colored in red and gray highlighted.

**Extended data 2.**
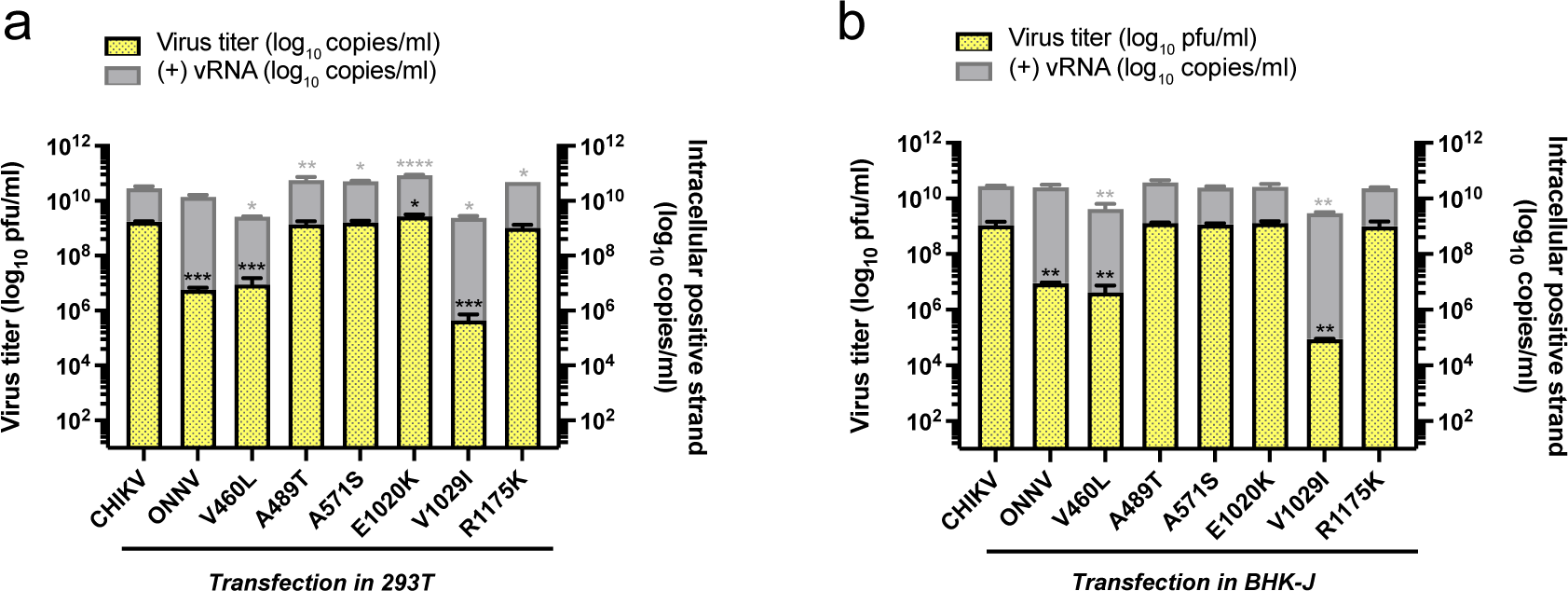
**a-b,** Infection of 293T **(a)** and BHK-J **(b)** cells with CHIKV vaccine strain 181/clone 25 positive selection site mutants. Viral replication and production of positive selection site mutants (V460L, A489T, A571S, E1020K, V1029I, R1175K) were determined by levels of intracellular (+) vRNAs and secreted infectious particles as previously described (One-way ANOVA and Dunnett’s multiple comparisons test: *, p≤0.05; **, p≤0.01; ***, p≤0.001; ****, p≤0.0001).

**Extended data 3.**
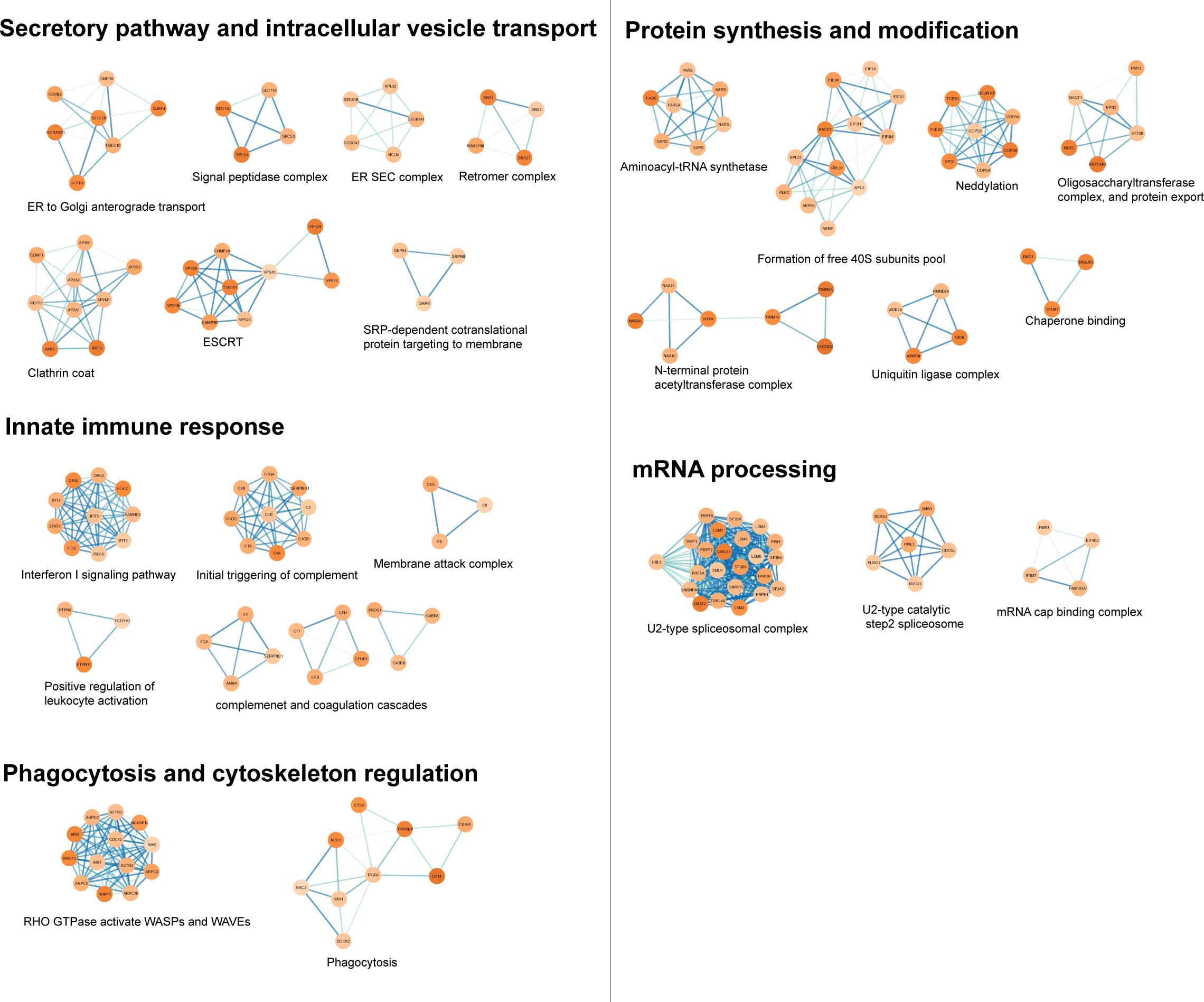
Enriched protein clusters from the host factors co-immunoprecipitated with CHIKV glycoproteins. The protein-protein interaction (PPI) of host factors detected in AP-MS were analyzed by STRING network and further clustered by MCODE^80^ and annotated with CORUM^88^ to classify functional protein clusters.

**Extended data 4.**
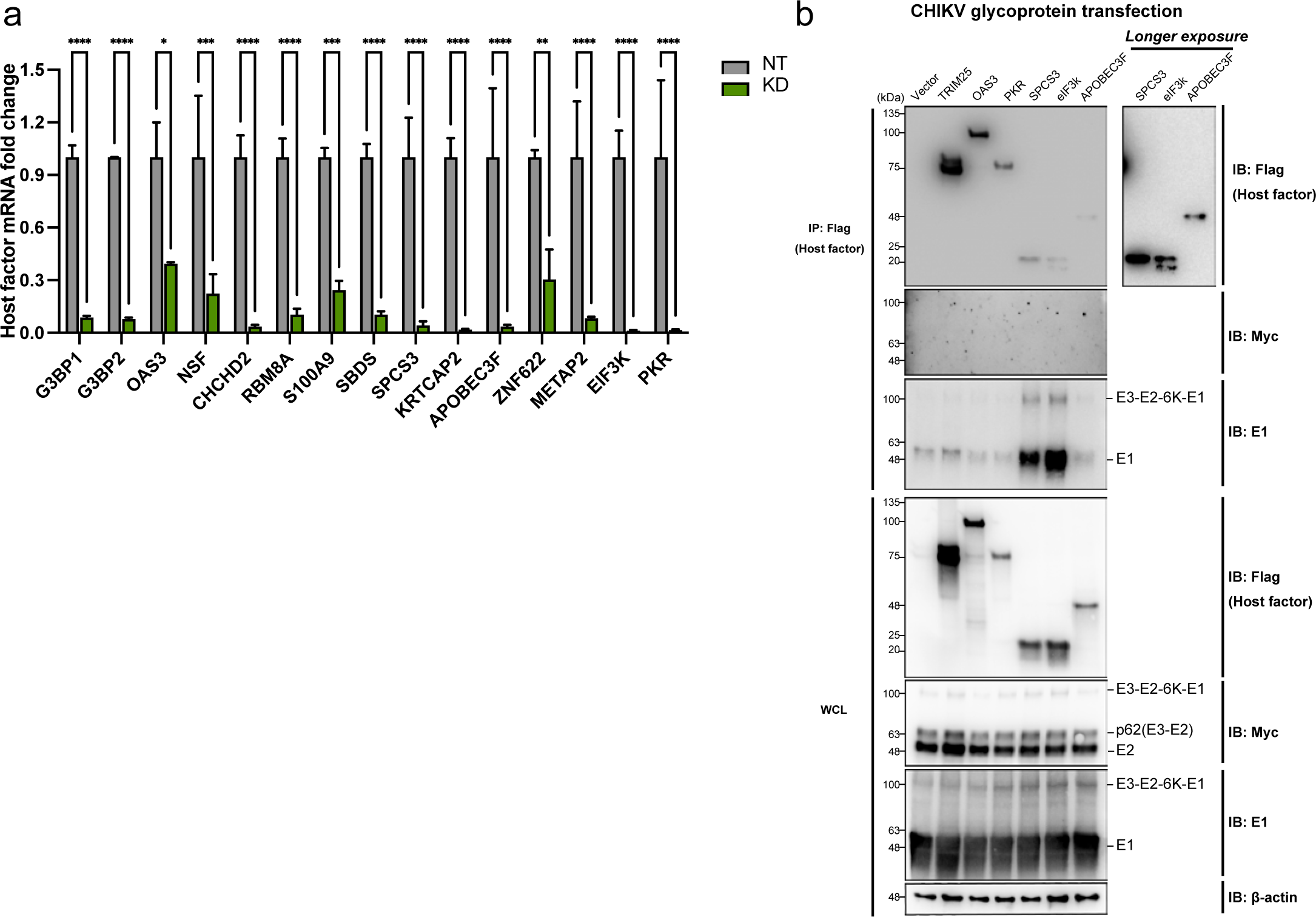
**a,** The macrophages were transfected with 25nM nontargeting siRNAs (NT) or pooled siRNAs targeting host factors (G3BP1, G3BP2, OAS3, NSF, CHCHD2, RBM8A, S100A9, SBDS, SPCS3, KRTCAP2, APOBEC3F, ZNF622, METAP2, EIF3K, and PKR). mRNAs of cells treated with siRNAs were extracted 48 h post transfection for RT-qPCR to evaluate the host factor knockdown efficiencies. (Two-way ANOVA and Šidák’s multiple comparisons test: *, p≤0.05; **, p≤0.01; ***, p≤0.001; ****, p≤0.0001). **b,** 293T cells were transfected with plasmids expressing 3xFLAG-taged host factors (TRIM25, OAS3, SPCS3, APOBEC3F, eIF3k, and PKR) or empty vector control for 24 h, and later transfected with plasmid expressing CHIKV glycoproteins (E3-myc-E2-6K-E1). The cells were lysed and immunoprecipitated by anti-FLAG agarose beads. Immunoblot was probed to check for E2/E1 binding to these host factors. TRIM25-3xFLAG was transfected into 293T cells for immunoprecipitation control. Data are representative of 3 independent experiments.

